# Preclinical efficacy of combinatorial B7-H3 CAR T cells and ONC206 against diffuse intrinsic pontine glioma

**DOI:** 10.1101/2025.09.01.673023

**Authors:** Andrea Timpanaro, Edward Z. Song, Ryma Toumi, Leonel Elena-Sanchez, Michael Meechan, Caroline Piccand, Kelsey Nemec, Anja Kordowski, Davina Lau, Scott Johnson, Lily Winter, Ashmitha Rajendran, Rebecca Ronsley, Shannon K. Oda, Joshua Gustafson, Jason P. Wendler, Carl Koschmann, Myron Evans, Siobhan Pattwell, Michael C. Jensen, Jessica B. Foster, Matthew D. Dun, Matthew C. Biery, Nicholas A. Vitanza

**Affiliations:** Ben Towne Center for Childhood Cancer and Blood Disorders Research, Seattle Children’s Research Institute, Seattle, WA, USA; Translational Cancer Research, Department for BioMedical Research (DBMR), University of Bern, Switzerland; Department of Pediatrics, Seattle Children’s Hospital, University of Washington, Seattle, WA, USA; Biomedical Informatics and Medical Education, University of Washington School of Medicine, Seattle, WA; Seattle Children’s Therapeutics, Seattle, WA, USA; Division of Pediatric Hematology/Oncology, Department of Pediatrics, University of Michigan, Ann Arbor, MI, USA; Division of Oncology, Children’s Hospital of Philadelphia, Philadelphia, PA, USA; Department of Pediatrics, University of Pennsylvania Perelman School of Medicine, Philadelphia, PA, USA; Cancer Signalling Research Group, School of Biomedical Sciences and Pharmacy, College of Health, Medicine and Wellbeing, University of Newcastle, Newcastle, New South Wales, Australia; Precision Medicine Research Program, Hunter Medical Research Institute, Newcastle, New South Wales, Australia; Paediatric Stream, Mark Hughes Foundation Centre for Brain Cancer Research, College of Health, Medicine and Wellbeing, Newcastle, New South Wales, Australia

**Keywords:** DIPG, immunotherapy, CAR T cell therapy, B7-H3, ONC206

## Abstract

**Background:** Diffuse intrinsic pontine glioma (DIPG) is a fatal pediatric brain tumor affecting over 300 children annually in the United States. Chimeric antigen receptor (CAR) T cells are a targeted immune effector cell therapy with substantial clinical benefit against hematologic cancers. Against CNS tumors, CAR T cells targeting B7-H3, a protein highly expressed on DIPG, have rapidly advanced from preclinical studies to clinical trials. BrainChild-03 (NCT04185038), a phase 1 trial of repeatedly delivered intracerebroventricular (ICV) B7-H3-targeting CAR T cells (B7-H3 CAR T cells), demonstrated tolerability and potential efficacy for children and young adults with DIPG. However, clinical benefits were not uniformly seen, and multi-agent treatment strategies may be required against such an aggressive disease. Here, we combined B7-H3 CAR T cells with ONC206, an imipridone molecule also under clinical investigation.

**Methods:** We tested B7-H3 CAR T cells combined with ONC206 across multiple DIPG cell cultures and orthotopic xenograft mouse models.

**Results:** B7-H3 CAR T cell monotherapy induced robust cytotoxicity while ONC206 treatment resulted in significant mitochondrial dysfunction against DMG/DIPG cells. The combination of low effector-to-target ratios of B7-H3 CAR T cells and IC_50_ concentrations of ONC206 led to significantly enhanced cytotoxicity *in vitro* (p<0.003) and increased IL-2, IL-29, VEGF-A, and Granzyme B levels. In vivo combinatorial studies of ONC206 and a single ICV dose of B7-H3 CAR T cells significantly extended survival in multiple DIPG xenograft mouse models (p<0.05).

**Conclusions:** B7-H3 CAR T cells combined with ONC206 is a feasible and efficacious multi-agent approach against multiple DIPG models.

**Importance of the study:** Diffuse intrinsic pontine glioma (DIPG) is a fatal pediatric brain tumor. While B7-H3 CAR T cells have shown tolerability and potential benefit in early trials, combinatorial regimens may be required for consistent cures against this aggressive disease. This study demonstrates that a preclinical therapeutic regimen of B7-H3 CAR T cells with ONC206, a second-generation imipridone, increases anti-tumor efficacy in vitro and in orthotopic DIPG mouse models. To our knowledge, this is the first study to evaluate ONC206 in combination with CAR T cells. Our findings provide a preclinical roadmap for evaluating small molecules with CAR T cells to interrogate both their combined benefit and the effect of small molecules on T cells themselves. This work offers a biologically-informed, clinically translatable strategy integrating small molecule therapeutics with CAR T cell therapy and support the development of multi-agent immunotherapy trials for children with DIPG and other high-grade brain and spinal cord tumors.

**Key Points:**

- B7-H3 CAR T cells are cytotoxic against preclinical DMG models.
- ONC206 causes metabolic apoptosis in preclinical DMG models.
- B7-H3 CAR T cells and ONC206 have combinatorial efficacy against DMG.

## Introduction

Diffuse intrinsic pontine glioma (DIPG), the most common form of diffuse glioma (DMG), is a fatal pediatric brain tumor most diagnosed in children between the ages of 5 and 10 years^1^. The prognosis remains dismal, with a median survival time of less than 12 months from diagnosis. The treatment of DIPG is challenging due to its critical location in the brainstem, where substantial resection is not feasible. These tumors further harbor resistance to conventional therapies underscoring a critical need for novel therapeutic approaches^2^. Recent advances in immunotherapy, particularly chimeric antigen receptor (CAR) T cell therapy, have opened new avenues for cancer treatment. B7-H3, a transmembrane protein involved in regulating T cell response^3,4^ has emerged as a promising therapeutic CAR T cell target on DIPG^5–8^. This cell surface protein was also found to be upregulated in a range of tumors including atypical teratoid rhabdoid tumors (ATRT)^9^, craniopharyngioma^10,11^, glioblastoma (GBM)^12,13^, high-grade glioma (HGG)^6,14^, medulloblastoma (MB)^15,16^, meningioma^17,18^, neuroblastoma (NB)^19^, and rhabdomyosarcoma (RMS)^20,21^.

Seattle Children’s first-in-human phase 1 clinical trial (BrainChild-03, NCT04185038) investigated safety and feasibility of B7-H3-targeting CAR T cells (B7-H3 CAR T cells) as locoregional immunotherapy against DIPG, DMG, and recurrent or refractory pediatric central nervous system (CNS) tumors. This trial demonstrated the tolerability of multiple intracerebroventricular (ICV) doses of B7-H3 CAR T cells at up to 100 million cells/dose given to young children in the outpatient setting^8^. While three patients were alive beyond four years from diagnosis, response to treatment was not uniform, suggesting the potential benefit of multi-agent therapy.

To enhance the efficacy of B7-H3 CAR T cells, we have investigated the combination with small molecules to modulate tumor susceptibility to immunotherapy. Among these, we tested imipridones, pro-apoptotic compounds characterized by a tricyclic core structure able to target dopamine receptor D2 (DRD2) and induce an anti-proliferative effect on cancer cells by modulating the AKT and ERK signaling pathways^22^. Novel studies have proven the capacity of imipridones to bind to the caseinolytic protease proteolytic subunit (ClpP) and activate the integrated stress response (ISR) cascade^23,24^. One class member, ONC201, has been evaluated in various tumor types including, brain cancers and demonstrated the ability to cross the blood-brain barrier^25,26^. ONC206, an analog of ONC201 with a greater binding affinity and selectivity^27^, follows the same mechanism of action by inducing apoptosis and stress-related signaling pathways in multiple models and may be more cytotoxic against DIPG^28^.

Here, we investigate the combinatorial potential of B7-H3 CAR T cells with ONC206 to enhance treatment efficacy against DIPG. Harnessing the tumor-targeting specificity of B7-H3 CAR T cells with the pro-apoptotic and stress-inducing properties of ONC206; this approach aims to increase tumor susceptibility to immunotherapy while maintaining treatment tolerability. Ultimately, this study explores the potential to combine targeted cellular therapy with small molecules to improve treatments against DIPG.

## Methods

All vendors and corresponding catalog numbers for the used materials are reported in Table 1.

**Table 1.** Material and methods list.

| ITEM | COMPANY | CATALOG NUMBERS |
| --- | --- | --- |
| **Cell culture** |  |  |
| NeuroCult™ NS-A Proliferation Kit (Human) | Stemcell Technologies | 5751 |
| RPMI-1640 | Gibco | 21870076 |
| DMEM | Gibco | 119600044 |
| Fetal Bovine Serum (FBS) | Gibco | A5670701 |
| Laminin | Trevigen | 3446-005-01 |
| Rat tail collagen type-I | Millipore Sigma | C3867 1VL |
| EGF | PeproTech | AF-100-15 |
| FGF | PeproTech | AF-100-18B |
| GlutaMax | ThermoFisher | 35050061 |
| Cryostor CS10 | StemCell Technologies | 100-1061 |
| P10 dishes | Greiner Bio-One CELLSTAR | 633181 |
| T75 flasks | Corning | 430641U |
| T150 flasks | Corning | 430825U |
| 96-well white flat bottom plates | Greiner Bio-One CELLSTAR | 07-000-138 |
| 96-well black flat bottom plates | Fisher Scientific | 08-772-225 |
| 24-well transparent flat bottom plates | Greiner bio-One | 662160 |
| D-Luciferin substrate | Revvity | 122799-5 |
| EasySep™ Human T cell isolation kit | StemCell Technologies | 17951 |
| Recombinant human IL 7 | BioLegend | 581904 |
| Recombinant human IL 15 | BioLegend | 570304 |
| **Plasmids** |  |  |
| psPAX2 | Addgene | 12260 |
| pVSV-G | Addgene | 8454 |
| **Caspase assays** |  |  |
| Incucyte Caspase 3/7 assay | Sartorius | 4440 |
| **Compounds** |  |  |
| Carboplatin (50 mg) | MedChem | HY-17393 |
| ONC206 | Chemspec API | 031000 |
| Etoposide | MedChem | HY-13629 |
| Everolimus (50 mg) | MedChem | HY-10218 |
| Marizomib (100 ug) | Sigma Aldrich | SML1916-100UG |
| Pacritinib | MedChem | HY-16379 |
| Paxalisib | MedChem | HY-19962 |
| Vincristine sulfate (10 mg) | SelleckChem | S1241 |
| Zotiraciclib | MedChem | HY-15163 |
| **Seahorse Technology** |  |  |
| Seahorse Cell Mito-Stress kit | Agilent Technologies | 103015-100 |
| XF RPMI medium, pH 7.4 | Agilent Technologies | 103576-100 |
| XF 1.0M Glucose | Agilent Technologies | 103577-100 |
| XF 100 mM Pyruvate | Agilent Technologies | 103578-100 |
| XF 200 mM Glutamine | Agilent Technologies | 103579-100 |
| CellTiter-Glo™ 2.0 Assay | Promega | PRG9243 |
| **Enzyme-linked immunosorbent assay** |  |  |
| IL-2 DuoSet ELISA development system | R&D Systems | DY202 |
| TNF-α DuoSet ELISA development system | R&D Systems | DY202-05 |
| IFN-γ DuoSet ELISA development system | R&D Systems | DY285B |
| **RNA extration** |  |  |
| RNeasy Plus Mini Kit | Qiagen | 74134 |
| **Primary Antibodies** |  |  |
| Brilliant Violet 421™ anti-human CD62L | BioLegend | 304828 |
| Brilliant Violet 510™ anti-human CD3 | BioLegend | 317332 |
| Brilliant Violet 605™ anti-human CCR7 | BioLegend | 353224 |
| FITC anti-human CD4 | BioLegend | 300530 |
| PECy7 anti-human CD95 | BioLegend | 305622 |
| PE anti-human CD45RA | BioLegend | 304108 |
| APC anti-human CCR5 | BioLegend | 359122 |
| Alexa Fluor® 700 anti-human CD8a | BioLegend | 301028 |
| PE anti-human B7-H3 | BioLegend | 351004 |
| FITC Annexin V Apoptosis Detection Kit | BioLegend | 640922 |

### Human specimen and patient-derived cell cultures

Human cell cultures were established with informed consent under institutional review board approval from Seattle Children’s Hospital (#14449). Biopsy specimens PBT-22FH, PBT-24FH, PBT-27FH, and PBT-29FH were acquired at diagnosis. The tumor tissues were procured at Seattle Children’s Hospital, and subsequent cell cultures were developed at Fred Hutchinson Cancer Research Center (FHCRC). SU-DIPG-XIII cell line was generously provided by Dr. Michelle Monje. These cell lines were cultured in NeuroCult NS-A Basal Medium supplemented with NS-A Proliferation Supplement (Stemcell Technologies), 40 ng/mL epidermal growth factor, and 40 ng/mL fibroblast growth factor (PeproTech), along with 2 mM GlutaMax (Gibco). Lenti-X 293T cells, used for CAR lentivirus production, were kindly provided by Prof. Michele Bernasconi and cultured in DMEM medium (Gibco) supplemented with 10% of fetal bovine serum (FBS) (Thermofisher). All cell culture models were validated with periodic DNA fingerprinting prior to experimentation.

### Lentivirus production

Lentiviruses (LVs) were generated via transfection of Lenti-X 293T cells, kindly provided by Prof. Bernasconi (University of Bern). 5x10^6^ 293T cells were seeded onto P10 plates, pre-coated with collagen type I rat tail, and cultured overnight in DMEM medium supplemented with 10% fetal bovine serum and 1x GlutaMax. The day after, 293T cells were co-transfected via calcium phosphate method with 10 µg of psPAX2, 4 µg of pVSV-G, and 10 µg of lentiviral B7-H3 CAR plasmid, as previously described^51^. Lentiviruses-containing supernatants were harvested 48 h post-transfection, filtered with 0.45 µm filters, and concentrated via overnight centrifugation at 4′000xg at 4 °C. Lentiviruses were aliquoted and stored at -80 °C until use.

### CAR T cell manufacturing

T cells were isolated from peripheral blood mononuclear cells (PBMCs) of healthy donors (Bloodworks Northwest) by negative selection using the EasySep™ human T cell isolation kit. T cells were then stimulated with anti-CD3/CD28 Dynabeads (Gibco, #40203D) at 1 × 10⁶ cells/mL for 36 h and transduced at a multiplicity of infection of 5 with LVs titered as previously described^51^. Transduced cells were counted every other day and maintained at a density of 0.8 × 10⁶ cells/mL in in RPMI-1640 medium supplemented with 10% fetal bovine serum, 1x GlutaMax, 5 ng/mL recombinant human IL-7, and 0.5 ng/mL recombinant human IL-15. After expansion, cells were cryopreserved in CryoStor® CS10 in liquid nitrogen until use.

### Bioluminescence survival assays

Bioluminescence killing assays were conducted by seeding at day 0 5’000 iRFP720^+^ firefly luciferase-positive (fLuc^+^) tumor cells onto a bioluminescence-compatible 96-well plate in 150 µl of NeuroCult medium (Stemcell). 50 µl of ONC206 at the final concentration of 150-300nM and/or 50 µl of B7-H3 CAR T cells at various effector-to-target (E:T) ratios of 1:10, 1:4, 1:2, 1:1, 2:1, and 5:1 were added to the respective wells at day 1. Untransduced mock T cells were used as negative controls. The other controls included 50 µl of medium replacing CAR T cells or ONC206. On the day of measurement, D-Luciferin was diluted in NeuroCult medium at the concentration of 30 µg/ml and administered to the wells. After 10-minute incubation at RT, bioluminescence intensity was quantified using a SpectraMax® iD3 Multi-Mode Microplate Reader (Danaher). Data from the assays were processed using Microsoft Excel and GraphPad Prism 9 for analysis.

### Live-imaging cytotoxicity assays

Two or seven-day-long cell viability assays were performed with live-imaging Incucyte technology to monitor tumor cell expansion and killing capacity of B7-H3 CAR T cells. 5’000 firefly luciferase^+^ (fLuc^+^) iRFP720^+^ tumor cells were plated at day 0 onto a black/clear bottom 96-well plate in 150 µl of NeuroCult medium. 50 µl of ONC206, at the final concentrations of 150-300nM, and/or 50 µl of B7-H3 CAR T cells at the various Effector-to-Target (E:T) ratios of 1:10, 1:4, 1:2, 1:1, and 5:1, were added to the respective wells at day 1. The respective control conditions included 50 µl of medium replacing CAR T cells and/or ONC206. Images were acquired every 4h using the Incucyte S3 device (Sartorius). Phase-contrast images were normalized to their respective baseline images taken at day 0. The results were processed with Incucyte 2022B Rev2 and Graphpad Prism 10 software.

### Enzyme-linked immunosorbent assays

50’000 PBT-22FH, PBT-24FH, PBT-27FH, PBT-29FH, and SU-DIPG-XIII cells were plated onto a 24-well plate and co-cultured with 250’000 B7-H3 CAR T cells (E:T ratio of 5:1) in 1ml of NeuroCult medium (Stemcell). After 24h, the supernatant was collected in 1.5ml tubes and stored at -80 °C until use. To assess the amount of human IL-2, TNF-a, and IFN-γ released by B7-H3 CAR T cells, ELISAs were performed following the manufacturer’s instructions. The results were analyzed with Microsoft Excel, whereas the figures were generated using GraphPad Prism 9.

### Flow Cytometry

Expression of surface targets was determined using the primary antibodies listed in Table 1. T cell phenotyping to determine the amount of naïve, central memory, and effector T cells included the expression of the following markers: CD62L, CD3, CD4, CD8, CD45RA. The expression of LAG-3, PD-1, and TIM-3 established exhaustion of CAR T cells before and after killing assays. Briefly, samples were washed twice in flow buffer (PBS supplemented with 2% BSA) and incubated at RT for 30 min with the corresponding primary antibodies at the dilution of 1:100. After washing cells twice, fluorescence was measured at the Cytoflex (Beckman). Flow analysis included a gate according to forward scatter/side scatter profiles and a gate on single cells. T cells were selected based on CD3 expression. Transduction efficiency was evaluated after 48 hours on EGFRt expression compared to untransduced T cells (Mock). FlowJo software was used to analyze flow cytometry data.

### Seahorse Assay

Oxygen consumption rate (OCR), basal respiration, non-mitochondrial respiration, and extracellular acidification rate (ECAR) were respiratory parameters measured using Seahorse XFe96 Analyzer and Cell Mito Stress Test Kit (Agilent). PBT-22FH, PBT-24FH, PBT-27FH, PBT-29FH, and SU-DIPG-XIII cells were treated with ONC206 at the different concentration of 150, 200, 250, and 300 nM. DMSO-treated cells were used as controls. After 48 hours, DIPG cells were washed and equilibrated to un-buffered medium for 1h at 37°C in a CO2-free incubator and transferred to the XFe96 analyzer. Reduction in mitochondrial respiration was assessed with sequential administration of 2 µM oligomycin, 2 µM carbonyl cyanide p-trifluoromethoxyphenylhydrazone (FCCP), and 0.5 µM Rotenone/ Antimycin A (AA). The experiments were performed following the manufacturer’s instructions, whereas the data were analyzed with Agilent Seahorse Wave Desktop software.

### Multiplex immunoassay

To evaluate cytokine release, 100’000 PBT-27FH and PBT-29FH cells were co-cultured with an equal number of CAR T cells in 24-well plates in the presence of ONC206 (250 nM) for 5 days in complete NeuroCult medium. Following incubation, supernatants were collected and analyzed using the human cytokine/chemokine 96-plex discovery assay® (HD96) (Eve Technologies, Calgary, AB, Canada). Cytokine concentrations were quantified and normalized as appropriate for subsequent analysis. All statistical analyses for the cytokine data were performed in R using the lme4 and emmeans packages. A single linear mixed-effects model was fitted, treating log2-transformed cytokine values as the response variable and including cell line, treatment, and cytokine as fixed effects; donor was included as a random intercept. Linear contrasts (e.g., “ONC206&CAR T vs. CAR T” and “ONC206 vs. PBS”) were calculated to assess differential expression in selected comparisons, with all p-values corrected using the false discovery rate.

### RNA sequencing

To perform RNA sequencing, 100,000 DIPG cells were co-incubated in 24-well plates with 100,000 CAR T cells in the presence of ONC206 (250 nM) for 5 days in complete NeuroCult medium. RNA was purified using RNeasy Plus Mini Kit (Qiagen) following the manufacturer’s instructions. RNA samples were sent to Novogene for sequencing. All pre-processing steps were run using Terminal. FastQ files were downloaded from Novogene and processed using FastP^52,53^ to filter reads and trim adaptors, followed by FastQC to evaluate quality. Reads were pseudoaligned to the human GRCh38 cDNA transcriptome (Ensembl release 113)^54^ using Kallisto (version 0.51.1)^55^ and run through MultiQC^56^ to assess post-mapping quality. All post-processing steps were run using RStudio (version 2024.12.0+467; R Core Team, 2022). Transcripts were summarized to the gene level via tximport (version 1.34.0)^57^ and abundance counts were calculated via TPM, followed by normalization using TMM via edgeR (version 4.4.2)^58–60^. PCA plots were made using the “prcomp” function. Differential gene expression analysis was performed using linear modeling via limma (version 3.62.2)^61^ and differentially expressed genes were called by applying cutoffs for FDR (0.05), counts per million (CPM) in at least one of the comparators (+1), and Log_2_(FC) (+/-2). Statistical overrepresentation tests and related GO Terms were calculated using the Panther Classification System (Version 19.0). Respective lists of upregulated DEGs were manually imported into the Gene List Analysis tab, selected for homo sapiens, and run through the statistical overrepresentation test option using the “GO biological process complete” option. Tests were run using the Fisher’s Exact option with an FDR threshold of 0.05. Related figures were created using GraphPad Prism (version 10.4.1).

### Surgical procedure and in vivo experimentation

8-week-old immunodeficient NOD.Cg-Prkdcscid Il2rgtm1Wjl/SzJ (NSG) female mice were purchased from Jackson Lab and housed at the Animal Facility of the Seattle Children’s Research Institute, under pathogen-free conditions. The experiments were approved by the Institutional Animal Care and Use Committee (IACUC) and conducted following the protocol #ACUC00669. Intracranial xenografts were established injecting 50’000 SU-DIPG-XIII cells, expressing a bicistronic cassette including mCherry and Firefly Luciferase (fLuc^+^), at 0.8 r lateral, 1.5 mm posterior, and 2.5 mm deep, relative to lambda.

Each cohort contained at least seven mice for adequate power. Only mice with confirmed tumor engraftment were maintained in the study and included in all analyses. 1 million Mock/CAR T cells were intracerebroventricularly (ICV) injected, whereas ONC206 was orally administered twice a week at the concentration of 50 mg/kg. For PBT-29FH study, little chunk tumors were chopped and injected using the same coordinates, whereas 250’000 Mock/CAR T cells were intracerebroventricularly (ICV) injected. Tumor growth was weekly monitored via optical imaging (In Vivo Imaging System), and mouse weights were measured twice a week. Symptomatic mice were euthanized, and tumors were post-mortem resected for IHC analysis.

### Statistical analyses

Statistical analyses were performed using GraphPad prism 9.0 (GraphPad Software, Inc., La Jolla, CA, USA). Dose-response curves were generated using nonlinear regression [log (inhibitor) vs. response - Variable slope (four parameters)]. Cell proliferation was analyzed using a two-way ANOVA (Sidak’s multiple comparison). Seahorse data were analyzed using one-way ANOVA (Tukey’s multiple comparison). For mice survival, Kaplan–Meier method (Kaplan–Meier survival fractions) was used to calculate the Log-rank (Mantel–Cox) and generate p-values. The statistical significance between experimental and control triplicate groups was analyzed incorporating a Dunnett’s multiple comparison test following a two-way ANOVA to examine main effects and interactions between variables. The levels of significance will be indicated as follows: p > 0.05 (ns), p ≤ 0.05 (*), p ≤ 0.01 (**), p ≤ 0.001 (***), p ≤ 0.0001 (****).

### Illustrations

Schemas and illustrations were generated with BioRender.com.

## Results

### On-target effector function of B7-H3 CAR T cells against DMG/DIPG models with high target expression

We first assessed the expression of B7-H3 on treatment-naïve, patient-derived DMG/DIPG models (PBT-22FH, PBT-24FH, PBT-27FH, PBT-29FH^29,30^) and an autopsy-derived model (SU-DIPG-XIII). Consistent with several other reports^5,6,31,32^, the quantification of surface copy numbers revealed a high density of B7-H3 at the protein level, with a minimum density of 30,000 molecules per cell and over 200,000 molecules on PBT-24FH (Figure 1A).

**Figure 1.**
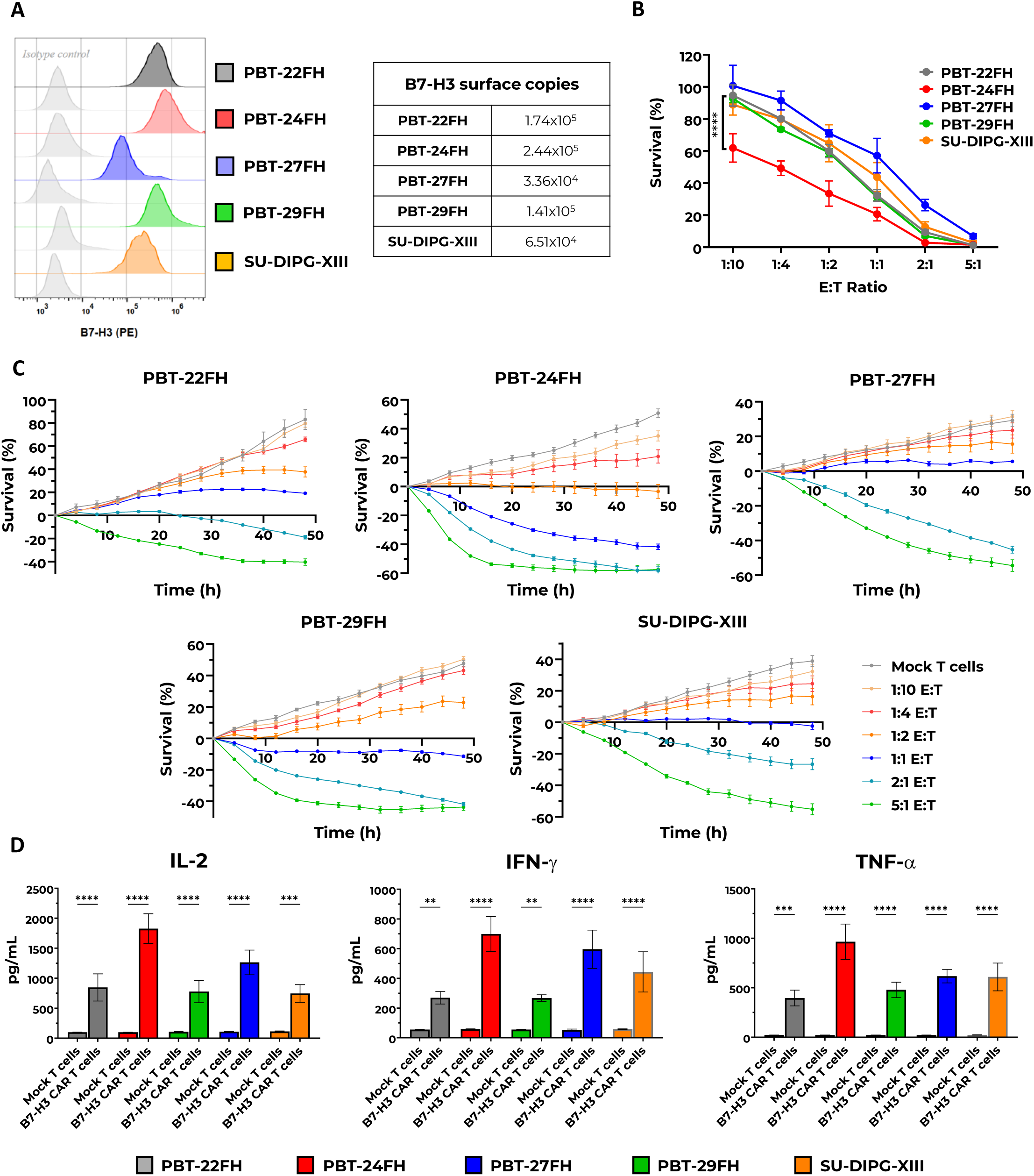
B7-H3 expression on DIPG models and efficacy of B7-H3 CAR T cells. **A.** B7-H3 expression was assessed in PBT-22FH, PBT-24FH, PBT-27FH, PBT-29FH, and SU-DIPG-XIII models using flow cytometry. High levels of B7-H3 were uniformly detected across all models with a PE-conjugated anti-B7-H3 antibody. Surface copy numbers of B7-H3 molecules were quantified with BD Quantibrite beads. **B.** B7-H3 CAR T cell killing capacity was determined after co-incubation of CAR T cells with fLuc^+^ iRFP720^+^ DIPG cells at different Effector-to-Target (E:T) ratios. Tumor cell survival (%) was evaluated at a single 48-hour time point using luciferase assays. Significantly higher killing was observed against PBT-24FH, compared to the other models. P<0.001 (n = 3 donors). **C.** Live-imaging experiments assessed the effectiveness of low-doses of B7-H3 CAR T cells (n = 3 donors) against DIPG models at multiple time-points, serving as a preliminary step to explore enhanced combinatorial killing with ONC206. These results, normalized to the respective measurements at t = 0h, report the most representative donor. **D.** Cytokine release of IL-2, IFN-γ, and TNF-α from B7-H3 CAR T cells was assessed via ELISA after 24h co-incubation with DIPG cells (n = 3 donors). ns = not significant; p < 0.05; p < 0.01; p < 0.001.

To assess B7-H3 CAR T cell efficacy, co-incubation experiments were performed at various effector-to-target (E:T) ratios (1:10, 1:4, 1:2, 1:1, 2:1, and 5:1) and cytotoxicity was evaluated after 48 hours. Data from three independent CAR T cell donors confirmed high killing ability, with >50% tumor cell lysis observed at the E:T ratio of 1:1 (Figure 1B). Notably, PBT-24FH - a hypermutant DIPG^33^ - exhibited greater sensitivity, with a 40% reduction in survival at the lowest E:T ratio of 1:10 (Figure 1B, red). To investigate long-term efficacy at low CAR T cell doses and to optimize conditions for evaluating potential synergy with ONC206, live-imaging experiments were performed considering multiple time-points (Figure 1C). Consistent with previous findings, an E:T ratio of 1:1 confirmed a cytostatic-like effect in most models, while PBT-24FH cells were eradicated more rapidly. Of note, higher doses of CAR T cells resulted in excessive tumor cell lysis, limiting the ability to distinguish potential synergistic effects of combination therapies. Therefore, E:T ratios of 1:10, 1:4, and 1:2 were selected for subsequent combinatorial studies with ONC206.

The efficacy of B7-H3 CAR T cells was further characterized by quantifying cytokine release, including IL-2, TNF-α, and IFN-γ. Supernatants were collected after 24 hours of co-incubation of CAR T cells with DIPG models. The results demonstrated that B7-H3 CAR T cells produced significantly elevated levels of IL-2, IFN-γ, and TNF-α compared to mock T cells, with the most robust cytokine response observed against PBT-24FH cells (∼1750 pg/ml IL-2, ∼1000 pg/ml IFN-γ, and ∼700 pg/ml TNF-α; Figure 1D).

These findings not only validated the cytotoxic activity of B7-H3 CAR T cells in our DIPG models but also guided the selection of optimal E:T ratios for subsequent studies investigating the combinatorial potential with ONC206.

### ONC206 demonstrates limited effects on CAR T cell proliferation

To explore promising therapeutic options to combine with B7-H3 CAR T cells, we selected several candidate drugs based on previously reported efficacy in preclinical and clinical DIPG models.

Marizomib has shown promise as a proteasome inhibitor capable of crossing the blood-brain barrier, warranting its inclusion in ongoing studies for DIPG treatment^34,35^. ONC206, an imipridone derivative of ONC201, has shown significant potency and improved efficacy against DIPG cells both *in vitro* and *in vivo*, making it a promising therapeutic candidate currently under clinical investigation^27,36^. Paxalisib, targeting the PI3K/mTOR pathway, has shown encouraging preclinical results and is being evaluated in clinical trials specifically for DMG/DIPG^37–39^. Zotiraciclib, a CDK9 inhibitor, exhibits strong anti-glioma effects in preclinical studies, suggesting its potential applicability in treating DIPG^40^. Additionally, everolimus, an mTOR inhibitor, and conventional chemotherapeutic agents, such as carboplatin, etoposide, and vincristine, were tested against pediatric CNS tumors.

To assess baseline drug sensitivity, IC_50_ values were determined after 120 hours of treatment using Cell Titer-Glo assays in two representative DMG models: the H3.1-mutant PBT-27FH and the H3.3-mutant PBT-29FH (Figure 2A-C). We then tested the drugs for potential compatibility with B7-H3 CAR T cell therapy. CAR T cells were co-cultured with tumor cells at a 5:1 E:T ratio for 24 hours to allow for antigen-mediated activation. Subsequently, each drug was added at its respective IC_50_ concentration, and CAR T cell survival and expansion were monitored for five days. The experimental workflow allowed for CAR T cell activation through antigen recognition before drug exposure, isolating the effect of the compounds on CAR T cell viability.

**Figure 2.**
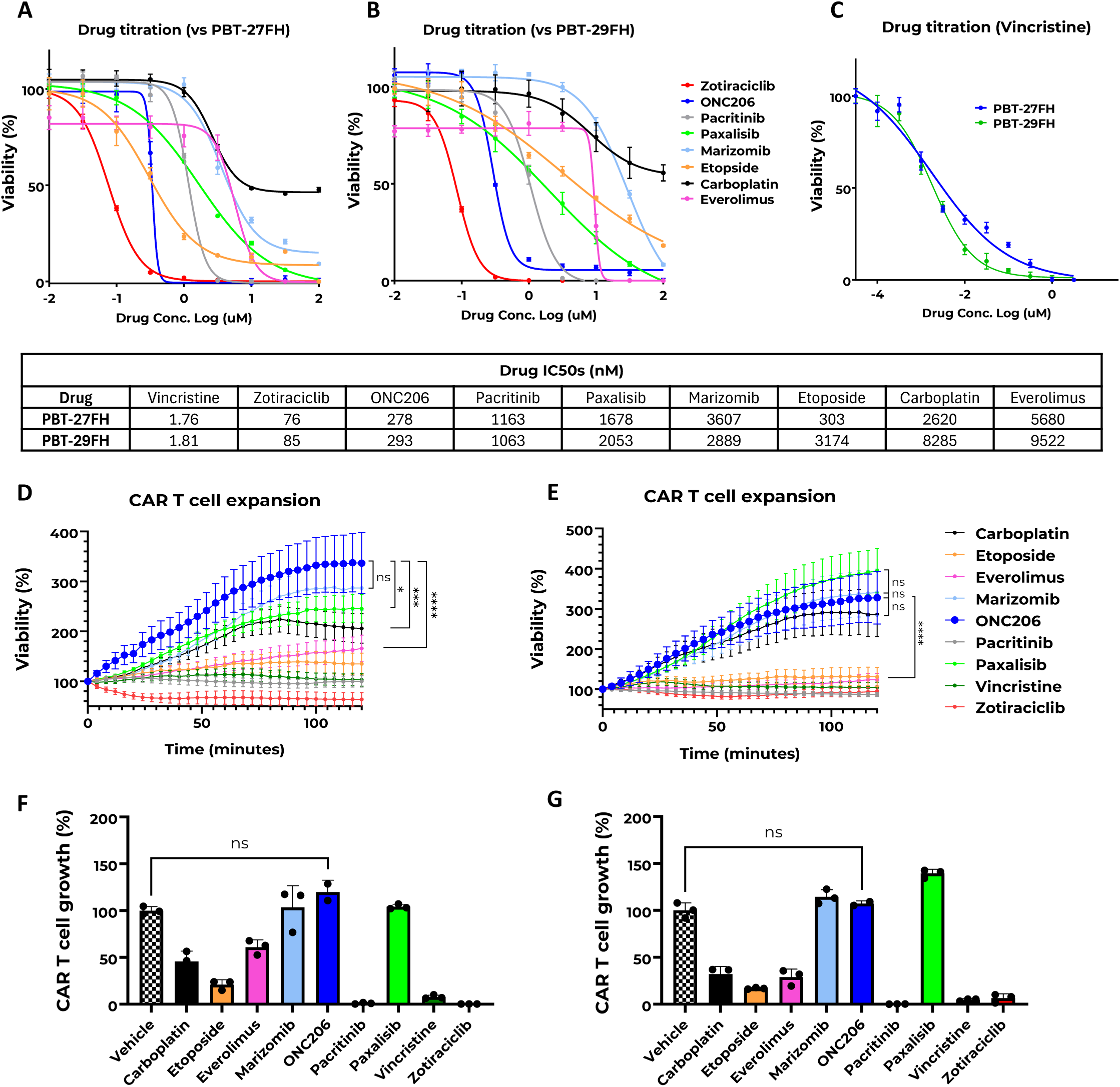
ONC206 selection from a 9-drug panel and its impact on B7-H3 CAR T cell viability and expansion. **A-B**. Dose-response curves of PBT-27FH (A) and PBT-29FH (B) DIPG cell lines following 120-hour treatment with a panel of drugs: Carboplatin, Etoposide, Everolimus, Marizomib, ONC206, Pacritinib, Paxalisib, Vincristine, and Zotiraciclib. IC₅₀ values were calculated for each condition. **C.** Vincristine titration curve shown using a log concentration range of –4.5 to 0.5 due to its high potency. IC₅₀ values were comparable between PBT-27FH and PBT-29FH cells. A summary of IC₅₀ values is provided in the accompanying table. **D-E.** B7-H3 CAR T cell expansion in co-culture with PBT-27FH (D) or PBT-29FH (E) cells over 120 hours. CAR T cells were co-cultured at an effector-to-target (E:T) ratio of 5:1 for 24 hours, followed by treatment with the respective drugs at their respective IC₅₀ concentrations. ONC206, Marizomib, Paxalisib, and Carboplatin did not impair CAR T cell expansion, whereas Etoposide, Everolimus, Pacritinib, Vincristine, and Zotiraciclib significantly inhibited CAR T cell proliferation. **F-G.** Quantification of CAR T cell growth at 120 hours post-drug exposure in PBT-27FH (F) and PBT-29FH (G) co-cultures, normalized to vehicle control. ns = not significant; *p* < 0.05; p < 0.01; *p* < 0.001.

As shown in the viability curves (pre-co-incubation with PBT-27FH - Figure 2D - or PBT-29FH cells - Figure 2E) and fold-change expansion compared to B7-H3 CAR T cell treated with vehicle (Figure 2F-G), ONC206 stood out for its dual profile: anti-tumor cytotoxicity and minimal adverse effects on CAR T cell expansion. In contrast, several other compounds either impaired CAR T cell proliferation or lacked comparable tumoricidal activity. Paxalisib was not further explored here due to a parallel project^41^, whereas marizomib and carboplatin, despite lack of toxicity on CAR T cells were discarded due to their low potency against DIPG cells. These findings led us to prioritize ONC206 for downstream combination strategies with B7-H3 CAR T cells.

### ONC206 demonstrates robust induced apoptosis across multiple DIPG models

ONC206, a member of the imipridone family, has emerged as a potential anti-cancer agent due to its abilities to specifically recognize ClpP and DRD2 and induce apoptosis through pathways involving mitochondrial dysfunction and stress response signaling^27,28,42,43^. To evaluate the cytotoxicity of ONC206, we performed dose titration experiments on five different DIPG/DMG models - PBT-22FH, PBT-24FH, PBT-27FH, PBT-29FH, and SU-DIPG-XIII - identifying an IC_50_ range of 217 - 298 nM after 120 hours (Figure 3A), leading to the selection of the averaged inhibitory dose of 250 nM for further studies.

**Figure 3.**
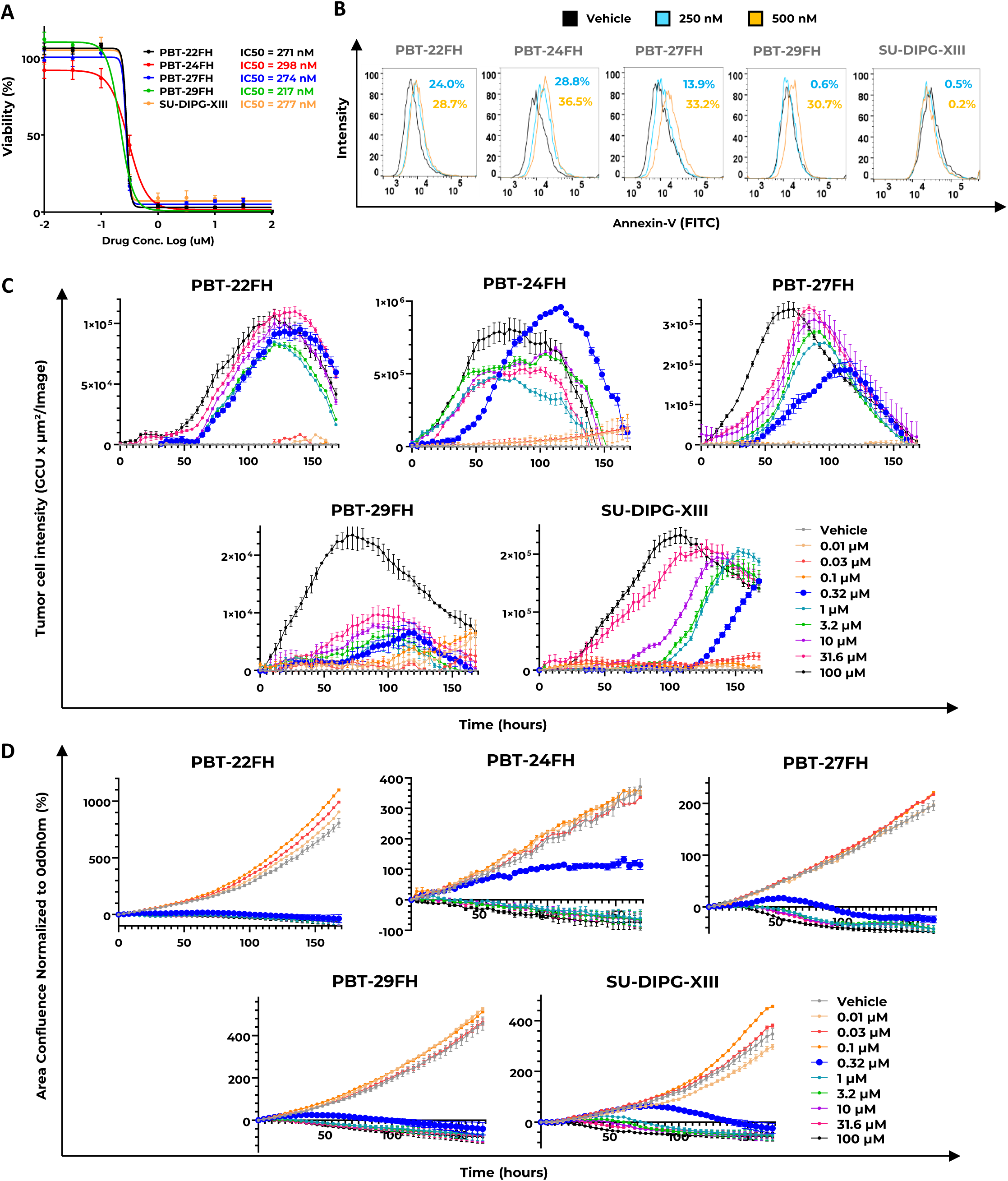
ONC206 cytotoxicity and apoptosis induction in DIPG models. **A.** ONC206 IC50 values were determined for PBT-22FH, PBT-24FH, PBT-27FH, PBT-29FH, and SU-DIPG-XIII cells. Tumor cells were incubated with varying concentrations of ONC206, and cell survival was assessed using cytotoxicity assays after 120 hours. Dose-response curves indicate consistent IC50 values ranging from 217 to 298 nM across DIPG cell lines. **B.** Early apoptosis was evaluated by flow cytometry. DIPG cells were treated with ONC206 at 250 nM (blue) and 500 nM (orange) for 48 hours, and Annexin-V levels were quantified using CytoFlex. **C.** Apoptosis induction by ONC206 was assessed in 7-day live-imaging assays by monitoring Caspase-3/-7 activation using a green fluorescence dye. An apoptotic peak near IC50 concentrations was observed at 120 hours in all models except SU-DIPG-XIII, which showed greater tolerance to apoptosis. **D.** Live-imaging experiments confirmed the efficacy of ONC206 at concentrations ≥0.32 µM. PBT-24FH and SU-DIPG-XIII exhibited partial resilience at 0.32 µM. All experiments were performed in biological triplicates (n = 3), with representative data shown.

Early apoptosis was evaluated using annexin-V staining and flow cytometry after 48 hours of treatment. For this experiment, a higher concentration of 500 nM was included to fully assess the apoptotic trend. As expected, a moderate dose-dependent increase in annexin-V levels was observed (Figure 3B), with higher responses at 500 nM, particularly in PBT-24FH, PBT-27FH, and PBT-29FH. In contrast, SU-DIPG-XIII showed minimal annexin-V expression (0.5% at 250 nM; 0.2% at 500 nM), indicating potential intrinsic resistance or activation of protective pathways in this model - similar ONC201 results were already observed in SU-DIPG-XIII model, revealing mechanisms linked to upregulated redox PI3K signaling^44^.

Live-cell imaging over seven days using a larger range of ONC206 concentrations (0.01 μM to 100 μM) and detecting caspase-3/7 expression via fluorescent dye confirmed a concentration-dependent apoptotic response (Figure 3C), where concentrations ≥1 μM induced rapid apoptosis in most models, while lower concentrations (<0.03 μM) had minimal effects. Remarkably, near-IC_50_ levels (0.32 μM) exhibited a distinct peak in caspase-3/7 activity at approximately 120 hours in most models, establishing this as optimal time-point for further experiments. As previously observed, SU-DIPG-XIII showed a delayed response with apoptosis peaking at 168 hours when ONC206 was used at 0.32 μM, supporting the notion of greater apoptosis-resistance. The inclusion of vehicle-treated controls further validated the specificity of the apoptotic response following negligible activation of caspase-3/7.

ONC206 caused significant cytotoxicity across all the cultures, with complete cell death observed within seven days at doses ≥0.32 μM (Figure 3D) highlighting effective cytotoxicity across the investigated DIPG models, with implications for combining it with B7-H3 CAR T cells to optimize therapeutic outcomes.

To further characterize the pro-apoptotic effects of ONC206 in DIPG, we performed RNA sequencing following a 120-hour co-culture with CAR T cells and ONC206 treatment (250 nM) against PBT-27FH and PBT-29FH. Sequencing yielded high-quality transcriptomes with Phred scores > 35 from two biological replicates per condition (Supplemental Fig.1A). Read alignment and mapping statistics confirmed efficient annotation to the human transcriptome (Supplemental Fig.1A). Principal component analysis (PCA) revealed distinct clustering between ONC206 and vehicle-treated samples, demonstrating strong treatment-driven transcriptional divergence in both models (Supplemental Fig.1B).

Differential expression analysis identified 6,953 genes significantly altered in PBT-27FH and 7,404 in PBT-29FH (adjusted p-value < 0.05). To refine our analysis to biologically meaningful transcripts, we applied stringent thresholds (Log_2_FC > 2 and CPM > 1), yielding 381 and 390 significantly upregulated genes in PBT-27FH and PBT-29FH, respectively (Supplemental Fig.1C). Comparison of these gene sets revealed a shared core of 140 commonly upregulated genes, with each model also exhibiting distinct transcriptomic signatures (Supplemental Fig.1D).

Gene ontology (GO) enrichment analysis of upregulated genes highlighted a strong overrepresentation of biological processes related to programmed cell death (GO:0043067), including subclasses such as positive regulation of apoptotic process (GO:0043065) and intrinsic apoptotic signaling pathway in response to endoplasmic reticulum stress (GO:0070059) (Supplemental Fig.1E). Building on our caspase-3/7 activation data, we focused specifically on the subset of pro-apoptotic genes within GO:0043065. A total of 37 pro-apoptotic genes were significantly upregulated in at least one of the two models upon ONC206 treatment, including key regulators such as ATF3, BBC3, BMF, DDIT3, FOSL1, and TNFRSF10B (DR5), supporting a consistent transcriptional shift toward apoptosis (Supplemental Fig.1F).

### ONC206 selectively disrupts tumor cell metabolism while preserving CAR T cell bioenergetics

Based on preliminary cytotoxicity results, we examined the effects of ONC206 on the metabolism of DIPG models at relevant concentrations near the IC_50_ (150 nM, 200 nM, 250 nM, and 300 nM). Given its known anti-tumor activity, we aimed to assess its impact on mitochondrial respiration and overall bioenergetic profiles, while also evaluating potential impairments in B7-H3 CAR T cell metabolism to determine its suitability for combination therapies. PBT-22FH, PBT-24FH, PBT-27FH, PBT-29FH, SU-DIPG-XIII, and CAR T cells were treated for 48 hours with ONC206, while control cells were cultured with the vehicle. After incubation, cellular metabolism was measured by using Seahorse technology, sequentially adding oligomycin, FCCP, and rotenone/antimycin A compounds to test mitochondrial stress.

Targeting cells at a low ONC206 dose (150 nM) demonstrated a selective impact on the metabolic profiles, highlighting a higher potential in inducing cell stress (Figure 4). Notably, a drastic dose-metabolic stress response was observed in all the DIPG lines tested in a dose-dependent manner (Figure S2). By contrast, CAR T cell metabolism was minimally impaired at IC_50_ doses (Figure 4A, bottom-right). In general, ONC206 treatment at 250 nM led to a complete collapse of mitochondrial respiration, as evidenced by the oxygen consumption rate (OCR) profiles (Figure 4A, red lines). Maximal respiration, as measured after FCCP treatment, was abolished, with tumor cells showing a lack of spare respiratory capacity (SRC) (Figure S2). Fold-change quantification across the DIPG cultures showed that ONC206 reduced basal respiration by over 90% compared to vehicles (Figure 4B, p<0.0001), indicating severe mitochondrial dysfunction. ATP-linked respiration and non-mitochondrial respiration were significantly reduced in most DIPG tumor cells (p<0.05 to p<0.001), except for PBT-24FH, which exhibited partial resistance (Figure S2). Additional respiratory parameters evaluated exhibited complete loss of SRC under the same conditions. Furthermore, bioenergetic profiles indicate that ONC206 effectively disrupts the respiratory capacity of DIPG cells, driving PBT-22FH, PBT-24FH, PBT-27FH, and PBT-29FH into a quiescent state, while shifting SU-DIPG-XIII toward a glycolytic phenotype (Figure 4C & Figure S2, right panels, for the other concentrations).

**Figure 4.**
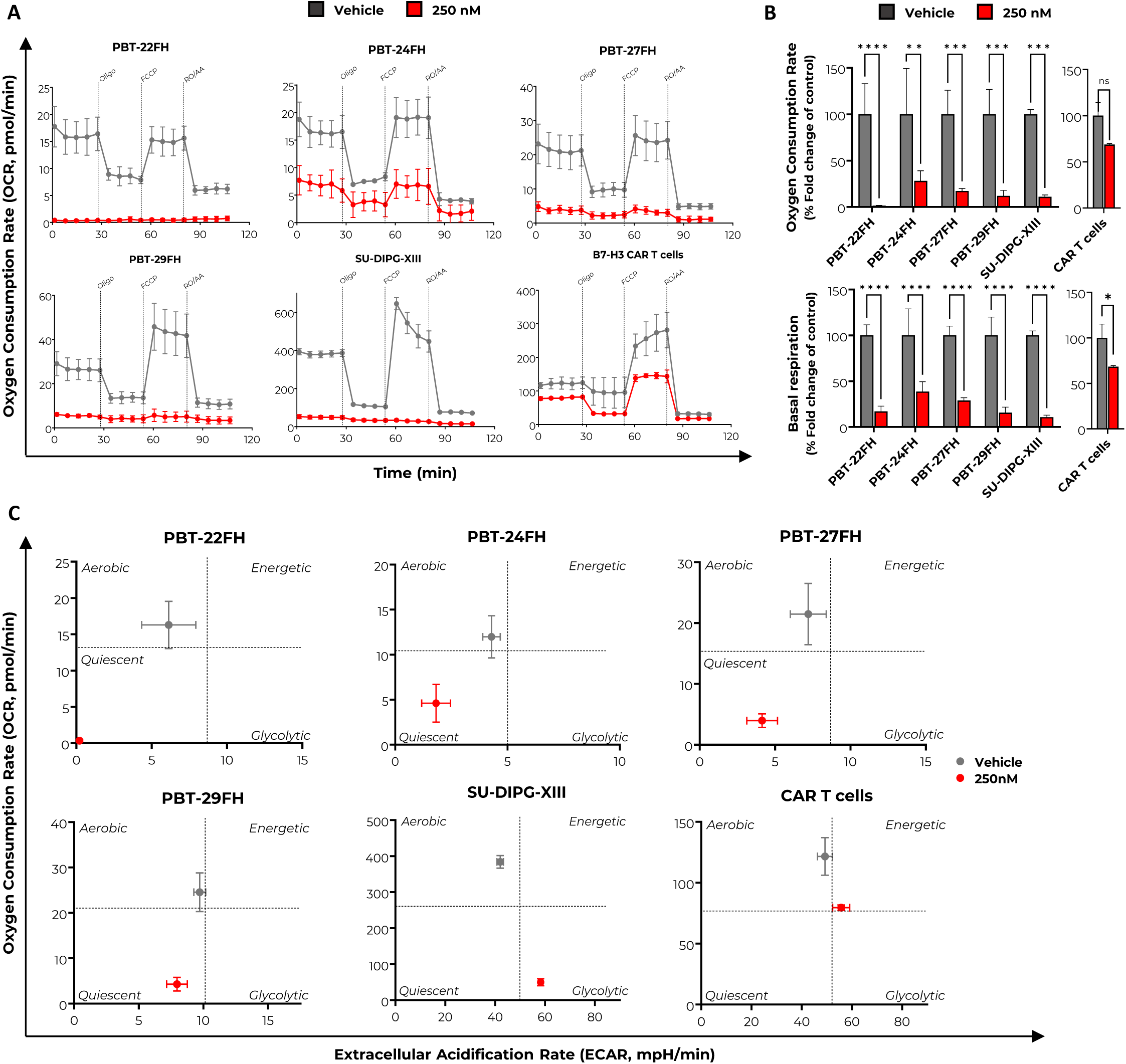
Respiratory response to ONC206 treatment in DIPG models and CAR T cells. PBT-22FH, PBT-24FH, PBT-27FH, PBT-29FH, SU-DIPG-XIII tumor cells and B7-H3 CAR T cells were treated with ONC206 at 250 nM (red) for 48 hours, and metabolic parameters were evaluated using Agilent Seahorse XF Technology. **A.** Oxygen consumption rate (OCR) profiles were measured over 100 minutes following sequential addition of oligomycin, FCCP, and rotenone/antimycin. Compared to the vehicle (gray), the IC50 dose (250 nM) impaired OCR in DIPG cells, while CAR T cells exhibited only a limited reduction. Fold-change bar charts for metabolic parameters, such as OCR and basal respiration, were quantified for all the investigated DIPG cells and CAR T cells, highlighting the significant resilience of CAR T cells at IC50 doses. **B.** Bioenergetics plots illustrate a metabolic shift in all the investigated DIPG models toward glycolytic or quiescent states under ONC206 treatment. In contrast, CAR T cells maintained an energetic phenotype, showing better tolerance to mitochondrial stress. All experiments were performed in triplicates (n = 3), with the most representative data shown.

On the other hand, CAR T cells displayed a different response to ONC206, underscoring their resilience to metabolic stress (Figure 4A, bottom-right panel). Despite a modest reduction in basal respiration (p<0.05), CAR T cells retained over 70% of their OCR under ONC206 treatment, reflecting preserved mitochondrial functionality (Figure 4B, lateral charts). ATP-linked respiration remained largely intact, with a negligible reduction compared to controls. Importantly, maximal respiration was maintained, with CAR T cells retaining their SRC under treatment conditions (p<0.05) (Figure S2X), whereas non-mitochondrial respiration decreased remaining functionally adequate (p<0.05) (Figure S2U). Notably, bioenergetic phenotype analyses underscored that CAR T cells preserved an energetic phenotype among all the experimental conditions with a well-preserved balance of OCR and extracellular acidification rate (ECAR) even at 300 nM (Figure S2Y). This ability to maintain metabolic flexibility and function despite ONC206 treatment highlights the remarkable resilience of CAR T cells to mitochondrial stress.

### Increased cytotoxicity of CAR T cell-mediated killing in combination with ONC206

Combinatorial experiments of ONC206 IC_50_ doses and low E:T ratios of B7-H3 CAR T cells were performed on PBT-22FH, PBT-24FH, PBT-27FH, PBT-29FH, and SU-DIPG-XIII to evaluate possible combinatorial benefits. DIPG cells were treated for 120 hours under optimal conditions as previously described for each cell line: Mock T cells (control), ONC206 (IC_50_-close concentrations), CAR T cells (E:T ratios in the range of 1:10 - 1:4), and combination. Killing capacity was evaluated via Incucyte experiments and luciferase assays.

Seven-day live-imaging experiments revealed a regulated combinatorial killing effect across the entire panel of models (Figure 5A). Single-agent treatments with CAR T cells (orange) and ONC206 (blue) primarily exhibited a cytostatic-like effect compared to the mock group (red). Notably, the impact of the combinatorial treatment (green) became increasingly pronounced over time, leading to statistically significant reduction in tumor cells at later time points, with greater tumor elimination by day 7 in PBT-22FH, PBT-27FH, and PBT-29FH models. Luciferase assays were further performed to measure tumor cell survival after 120 hours and corroborate these results. D-Luciferin was added at the concentration of 10 µg/mL into the wells, and tumor cell viability was normalized to the vehicle controls. Single treatments showed an efficacy of around 50% in each cell line, while combinatorial groups (green) revealed significantly superior killing across the DIPG culture panel (Figure 5B). These results, confirmed among three different CAR T cell donors, support an improved combinatorial effect mediated by ONC206, which is potentially able to sensitize tumor cell to immune-mediated cytotoxicity, offering the rationale for further preclinical studies.

**Figure 5.**
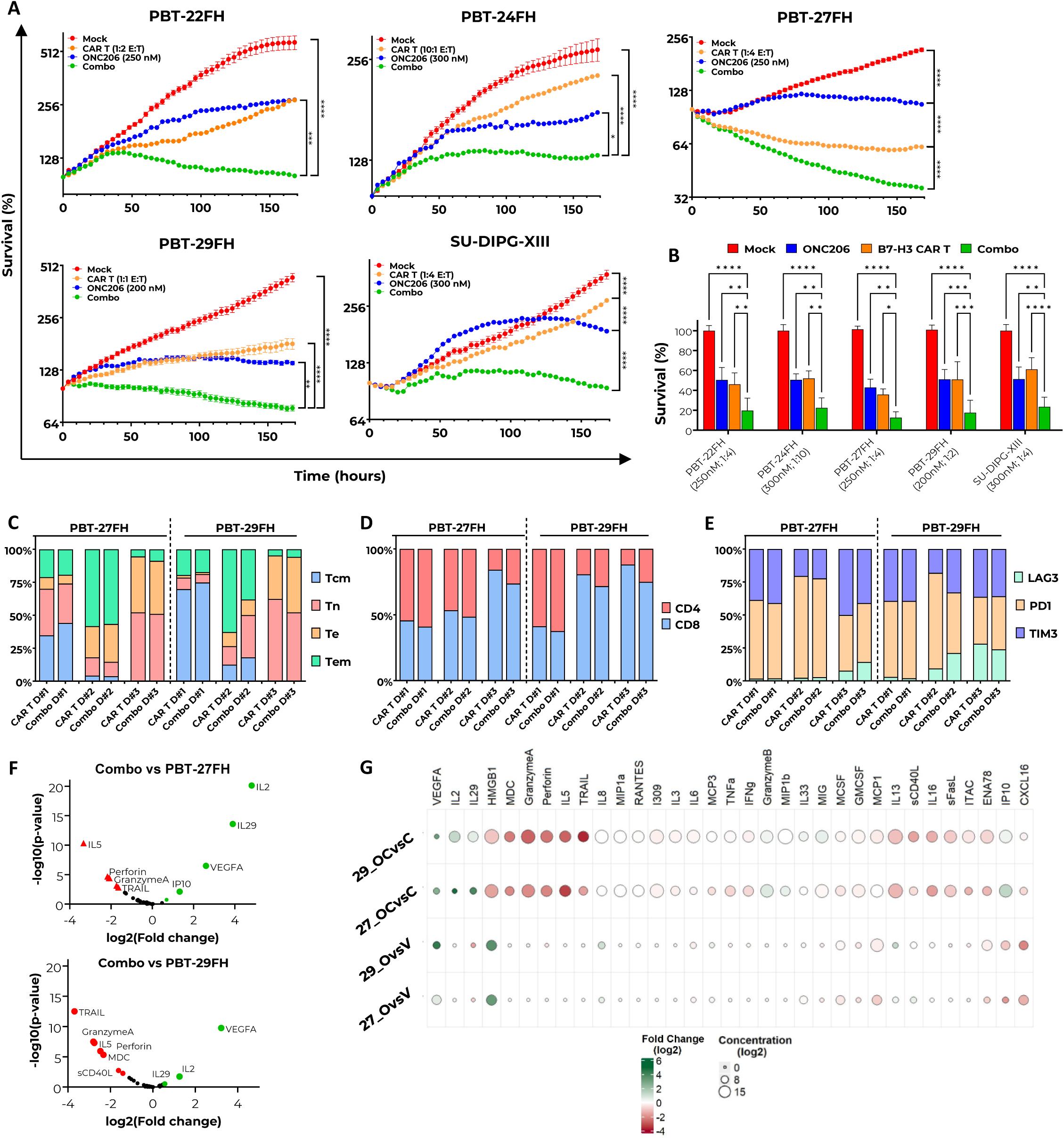
Low doses of ONC206 improve killing and preserve functional properties of B7-H3 CAR T cells. DIPG cell killing was evaluated under combinatorial conditions of ONC206 and B7-H3 CAR T cells, while variation of CAR T cell properties was assessed via flow cytometry and multiplex immunoassays after 5 days of ONC206 treatment. **A.** Incucyte technology was used to monitor DIPG cell survival for seven days. Combinatorial benefit (green) is compared to single treatments of low E:T ratios of CAR T cells (orange) and IC50 doses of ONC206 (blue). Mock T cells were used as negative control (red). **B.** Bioluminescence assays confirmed significant combinatorial benefit (green) after 120 hours of co-culture. **C-E.** CAR T cell phenotype, CD4/CD8 ratios, and CAR T cell exhaustion were evaluated by flow cytometry revealing non-significant changes between treated- and untreated-CAR T cells. D#1, #2, #3 represent the three T cell donors used. **F.** P-value vs fold-change charts highlight significantly increased levels of IL-2, IL-29, IP10, and VEGF-A, and decreased levels of TRAIL, Granzyme-A, IL-5, and perforins in PBT-27FH and PBT-29FH post-5-day treatment with ONC206 (250 nM). **G.** Bubble heatmap of 33 cytokines show contrasts of interest (“Combo vs CAR T alone” and “ONC206 vs vehicle”) for each cell line. Each bubble color indicates the log2 FC derived from the linear mixed-effects model (green = positive FC; red = negative FC). The bubble size corresponds to the average log2 cytokine concentration for each comparison’s control (no drug). All pairwise contrasts were FDR-adjusted.

### Functional profile of B7-H3 CAR T cells preserved at low doses of ONC206

To evaluate whether ONC206 IC_50_ dose influences CAR T cell effector functions, we analyzed T cell phenotypes by assessing the ratios of central memory (T_CM_), naïve-like (T_N_), effector (T_E_), and effector memory (T_EM_) T cells across three different donors. B7-H3 CAR T cells were co-incubated with the representative models PBT-27FH and PBT-29FH at the E:T ratio of 5:1, either in the presence or absence of ONC206 (250 nM). After 5 days, CAR T cells were collected and analyzed by flow cytometry. CAR T cells were then gated for CD3 expression, and T cell subtypes were determined based on CD62L and CD45RA expression. Upon antigen stimulation by either tumor cell model, the proportions of T_CM_ (CD62L^+^/CD45RA^-^), T_E_ (CD62L^-^/CD45RA^+^), and T_EM_ (CD62L^-^/CD45RA^-^) populations remained consistent (Figure 5C), despite visible variability among donors.

Similarly, the CD4/CD8 ratio in the combination groups was unchanged, compared to the respective controls (Figure 5D).

Next, we evaluated the expression of the markers LAG-3, TIM-3, and PD-1 to determine whether ONC206 affects CAR T cell exhaustion. Consistent with the previous results, no differences were observed in the percentage of CAR T cells expressing these markers between the controls and combination groups (Figure 5E) indicating that low doses of ONC206 do not significantly alter CAR T cell phenotypic composition, or CD4/CD8 ratios, nor do they cause CAR T cell exhaustion under these conditions.

### ONC206 IC_50_ dose does not impair cytokine release in activated CAR T cells

To assess the impact of ONC206 on cytokine secretion by B7-H3 CAR T cells, we conducted a comprehensive 96-plex immunoassay, co-culturing CAR T cells with PBT-27FH or PBT-29FH cells at an effector-to-target (E:T) ratio of 1:1 in the presence or absence of ONC206 (250 nM). Following five days of co-culture, supernatants were collected, and cytokine concentrations were quantified and normalized to vehicle-treated controls.

Linear contrast analyses from three independent T cell donors (e.g., fold change comparisons referred to as “ONC206 & CAR T vs. CAR T” and “ONC206 vs. PBS”) revealed that ONC206 does not globally suppress cytokine release by CAR T cells (Figure 5). Regardless of the DIPG model employed to stimulate CAR T cell activation, exposure to ONC206 resulted in a reduction of specific cytokines, including TRAIL, Granzyme A, IL-5, and perforin, compared to controls (Figure 5F, red dots). This effect however was counterbalanced by the significant upregulation of key cytokines such as IL-2, IL-29, IP-10, and VEGF-A (Figure 4F, green dots). Importantly, cytokines typically associated with CAR T cell characterization, including IL-3, IL-6, IL-8, TNF-α, IFN-γ, and Granzyme B, remained consistently sustained even after five days of ONC206 treatment, predominantly around a log2 value of 10, as visualized in the quantitative bubble heatmap (Figure 5G, white circles).

These findings demonstrate that ONC206 at 250 nM preserves CAR T cell cytokine secretion while selectively modulating specific pathways, thereby maintaining overall immune functionality. The sustained production of key cytokines, together with the targeted enhancement of others, supports its profile as a well-tolerated agent that maintains CAR T cell function and therapeutic potential. These results further support the rationale for combining ONC206 with B7-H3 CAR T cells, as its modulatory effects on T cells are likely minimal.

### Combination therapy with ONC206 and B7-H3 CAR T cells improves survival and maintains tolerability in orthotopic DIPG mouse models

After demonstrating that low doses of ONC206 significantly increased the killing capacity of B7-H3 CAR T cells *in vitro* without impairing T cell phenotype, exhaustion, and release of functional cytokines, we tested the proposed combinatorial approach in two distinct DIPG mouse models: SU-DIPG-XIII and PBT-29FH. For the SU-DIPG-XIII model, 5x10^4^ firefly luciferase-expressing tumor cells were stereotactically injected into NSG mice on day 0 (Figure 6A). Following tumor engraftment and normalization, mice were stratified into four experimental cohorts: a) Mock T cells + vehicle (control); b) Mock T cells + ONC206 (ONC206 monotherapy); c) B7-H3 CAR T cells + vehicle (CAR T monotherapy); and d) B7-H3 CAR T cells + ONC206 (combination therapy). ONC206 administration began on day 3, with a dosing regimen of 50 mg/kg, once daily, twice weekly, maintained until the experiment conclusion. At day 7, 1x10^6^ Mock T cells or CAR T cells were ICV delivered. Mice were monitored daily for potential treatment-related side effects, while body weight was recorded weekly. Animals were euthanized upon reaching endpoint criteria, including severe neurological symptoms or >20% body weight loss. The results demonstrated that ONC206 at 50 mg/kg was well tolerated, with no observed side effects, confirming its suitability for *in vivo* administration. However, ONC206 monotherapy failed to show efficacy in the SU-DIPG-XIII model, as survival was comparable to the control group (Figure 6B). In fact, both the control and ONC206 monotherapy cohorts reached endpoint within 12 days post-T cell injection, with median survival times of 15 and 14 days, respectively. In contrast, the CAR T monotherapy group exhibited a substantial benefit, extending the survival of 2 out of 12 mice up to day 40, with a median survival of 25 days (Figure 6B-D). Notably, the combination therapy significantly outperformed both monotherapy groups (p < 0.05), demonstrating a clear advantage in delaying tumor progression as early as day 19. Remarkably, 6 out of 12 mice in the combination group survived beyond day 35 (median survival), and three mice achieved long-term survival, with one reaching up to 101 days (Figures 6B-D). Weight progression data further support the tolerability of the combination treatment, with weight loss occurring only at advanced stages of disease when tumor burden was high (Figure 6D).

**Figure 6.**
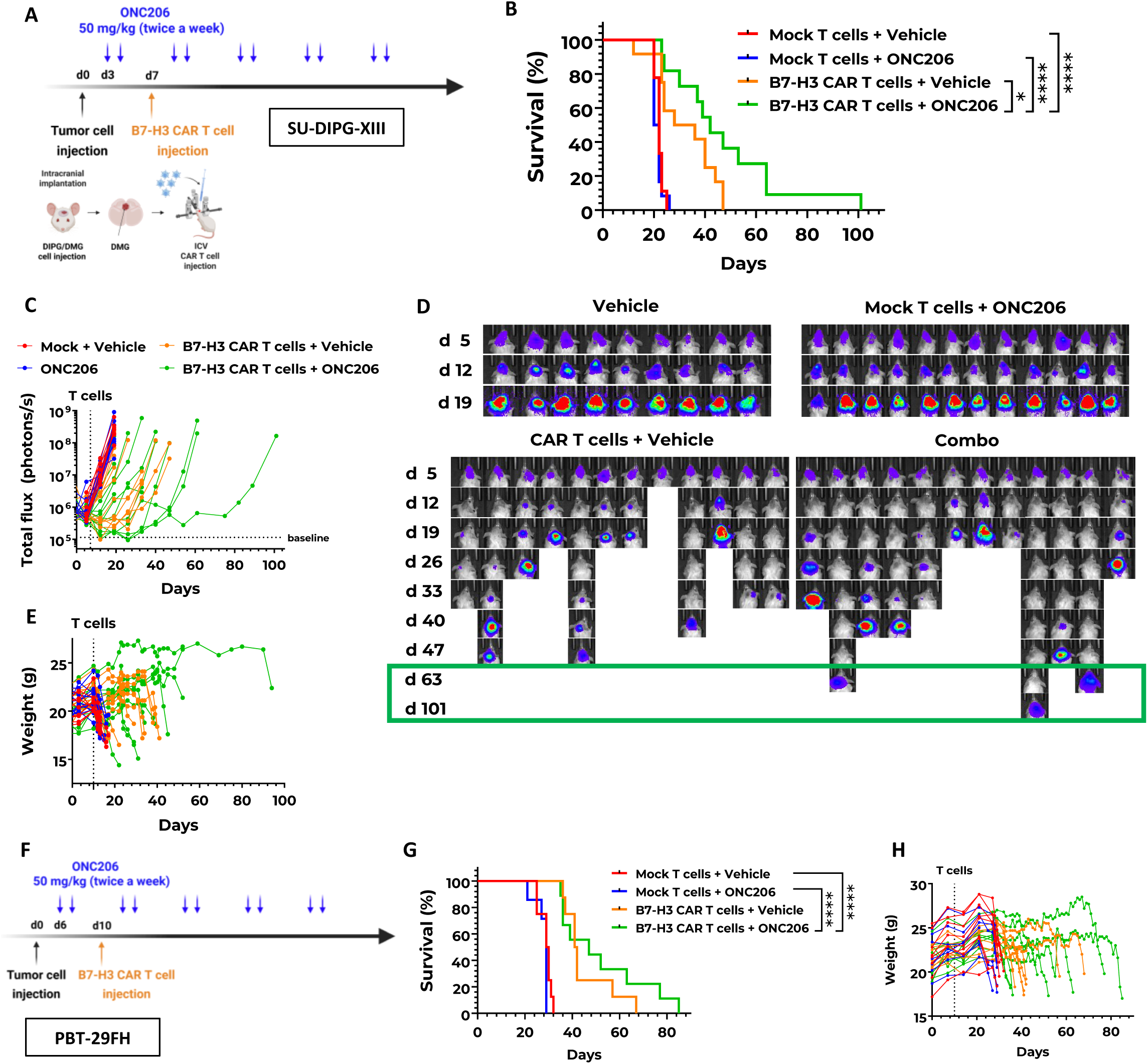
ONC206/B7-H3 CAR T cell combinatorial treatment showed significant extended survival in DIPG models. **(A)** Schematic representation of the experimental timeline for B7-H3 CAR T cell treatment in combination with ONC206 against the SU-DIPG-XIII model. Tumor-bearing mice received ONC206 (50 mg/kg) at the indicated time points following T cell injection. 1 million mock T cells/B7-H3 CAR T cells were ICV injected at day 7. **(B)** Kaplan-Meier survival curve comparing the different treated groups shows a subset of mice with long-term tumor control able to delay survival up to 94 days. **(C)** Tumor burden progression, measured by bioluminescence imaging (total flux), showing improved tumor control in the CAR T cell + ONC206 combination group. **(D)** Representative bioluminescence images of tumor burden over time for each treatment group. The CAR T cell + ONC206 combination resulted in the most effective tumor suppression. **(E)** Weight progression over time across treatment groups. No significant weight loss was observed, indicating treatment tolerance. **(F)** Schematic representation of the experimental timeline for B7-H3 CAR T cells treatment in combination with ONC206 against PBT-29FH model. **(G)** Kaplan-Meier survival curve shows extended survival up to 75 days in the combinatorial group compared to the B7-H3 CAR T cell treatment. Significant difference was observed with ONC206 monotherapy and mock group T cells. **(H)** Weight progression over time across treatment groups. (****p < 0.0001, *p < 0.05, log-rank test).

For the PBT-29FH model, murine tumor tissues were mechanically dissociated from mouse donors, and small fragments were uniformly implanted into NSG mice on day 0. Unlike the SU-DIPG-XIII model, bioluminescence imaging for PBT-29FH was not feasible, as the implanted tumor fragments lacked a luciferase reporter. Consequently, tumor progression was assessed solely based on mouse survival and weight monitoring. As in the SU-DIPG-XIII model, four experimental cohorts were established, maintaining the same treatment conditions. ONC206 was administered at 50 mg/kg, once daily, twice weekly, whereas 0.25 x10^6^ mock T cells or B7-H3 CAR T cells were ICV delivered at day 10 (Figure 6F). The Kaplan-Meier survival analysis (Figure 6G) demonstrated a significant survival benefit in the combination therapy group (green line) compared to the mice receiving control treatment (red) or ONC206 monotherapy (blue), the latter of which exhibited the shortest survival, indicating a limited therapeutic efficacy of ONC206 alone in this aggressive model. While B7-H3 CAR T cell monotherapy (orange) extended survival relative to controls, the combination therapy resulted in noticeable improvement, with survival extended until day 85. No toxicity was observed in the combination group, and mouse weight loss was related to tumor (Figure 6G, 6H). Together, the absence of neurological side effects and longer survival in the combination cohort during the whole treatment indicate that ONC206 at 50 mg/kg is well tolerated and does not contribute to significant adverse effects. Collectively, these results highlight that ONC206 may sensitize tumor cells and enhance B7-H3 CAR T cell-mediated tumor control, leading to significant survival benefit.

## Discussion

B7-H3 CAR T cell therapy has emerged as a promising strategy for treating DIPG, offering tumor selectivity, therapeutic tolerability, and potential clinical activity^7,8^. However, obstacles including tumor heterogeneity, a hostile tumor immune microenvironment, low CAR T cell persistence, and the need to target a diffusely invading disease often hamper effective treatment. Therefore, here we attempted to combine B7-H3 CAR T cells with other agents undergoing clinical advancement for improved efficacy.

Here, we confirmed that ONC206 robustly induces apoptosis and metabolic collapse in multiple DIPG models, while preserving the metabolic fitness, phenotype, and effector functions of B7-H3 CAR T cells. Seahorse analyses confirmed that, even at IC_50_ concentrations (150 - 300 nM), ONC206 selectively impairs tumor cell oxidative metabolism while sparing CAR T cell basal and maximal respiratory capacity. Furthermore, IC_50_-doses of ONC206 did not significantly alter T cell subset distribution, exhaustion marker expression, or CD4/CD8 ratios, even after prolonged co-culture. Cytokine profiling revealed preserved - or even enhanced - secretion of key effector molecules such as IL-2, VEGF-A, IL-29, IP10, IFN-γ, Granzyme B, and TNF-α by CAR T cells in the presence of ONC206, supporting the notion that IC_50_-doses of this compound do not compromise T cell functionality.

Critically, combination therapy with ONC206 and B7-H3 CAR T cells led to significantly improved tumor cell elimination and increased killing efficacy *in vitro* across multiple DIPG cultures via both luciferase assays and live-imaging experiments. Beyond *in vitro* analyses, the combinatorial efficacy was further supported *in vivo*, in which mice treated with low doses of B7-H3 CAR T cells and ONC206 exhibited markedly improved tumor control and had significantly extended survival in two distinct orthotopic DMG models. Notably, the combination resulted in the most extended survival observed, without treatment-related toxicity. This enhanced efficacy is likely mediated by mitochondrial stress induced by ONC206 and apoptotic priming in tumor cells, increasing their susceptibility to immune-mediated killing. However, despite a clear trend observed in our DMG/DIPG models, not all responded equally, underscoring the complexity of DIPG biology and the need for model-specific strategies.

Limitations of this work include the lack of repeated intracranial CAR T cell dosing, which is a primary strategy currently used in patients. As CNS catheter implantation systems have emerged^45,46^, future studies will incorporate less aggressive *in vivo* models to provide an opportunity for long term ONC206 dosing and repeated ICV CAR T cell doses. This work will allow future longer-term studies that have the potential to allow for blocks of therapy and other innovative combinatorial strategies. Further, we are yet to incorporate immunocompetent models, though work is currently ongoing to pilot recently developed DIPG syngeneic models and mouse-specific B7-H3 CAR T cells. These models will provide avenues for assessments of potential immune modulation by ONC206 and of potential inhibitory microenvironmental factors that must be addressed for patient therapeutic success. Importantly, our data align with emerging insights from Ziegler and colleagues^47^, who recently demonstrated that the sequence in which combination therapies are delivered can critically influence treatment efficacy in DIPG. This reinforces the rationale for building strategically ordered regimens that both sensitize tumors and support immune function.

Ultimately, this work contributes to a growing, joint international effort to develop rational, mechanism-informed combination approaches that have a higher chance of translational success^48–50^. Future studies should define the optimal sequence and timing for integrating B7-H3 CAR T cells with ONC206 and exploring its potential in other B7-H3-expressing or immune-evasive tumors.

## Data availability statement

All requests for raw and analyzed data and materials will be promptly reviewed by the Intellectual Property Office of Seattle Children’s Research Institute to determine whether the request is subject to any confidentiality or intellectual property restrictions. Raw preclinical data are securely stored at Seattle Children’s with indefinite backup. The data supporting the findings of this study are available from the corresponding author upon reasonable request. Any data and materials that can be shared via a Material Transfer Agreement.

## Acknowledgements

The authors would like to acknowledge the generosity of all patients and their families. We thank the services provided by Seattle Children’s administrative staff, Flow Cytometry Core, and Office of Animal Care, as well as the Experimental Histopathology Shared Resource (RRID:SCR 022606) of the Fred Hutch/University of Washington/Seattle Children’s Cancer Consortium (P30 CA015704). We thank Chimerix for providing the drug ONC206, although Oncoceutics, Chimerix, and Jazz Pharmaceuticals did not provide salary support or research funding for any component of this project.

## Authorship statement

AT: Design and execution of experiments, supervision, analysis, interpretation of data, writing, and funding acquisition. EZS: Design and execution of experiments, analysis and interpretation of data. RT: Design and execution of experiments, analysis and interpretation of data. LES: Execution of experiments, analysis and interpretation of data. MM: Execution of experiments, analysis and interpretation of data. CP: Execution of experiments, analysis and interpretation of data. KN: Execution of analysis and interpretation of data. AK: Execution of experiments, analysis and interpretation of data. DL: Execution of experiments. SJ: Execution of experiments. AR: Execution of analysis and interpretation of data. JPW: Execution of experiments, analysis and interpretation of data. RR: critical reading of the manuscript. JG: Resources and methodology. CK: Critical reading of the manuscript. ME: Supervision. SP: Supervision. MCJ: Supervision, resources, and critical reading of the manuscript. JBF: Resources, methodology, and critical reading of the manuscript. MDD: Resources and methodology. MCB: Supervision and critical reading of the manuscript. NAV: Design of the study, interpretation of data, supervision, writing, and funding acquisition. All authors read and approved the final manuscript.

