## Supplemental Figures for "Preclinical efficacy of combinatorial B7-H3 CAR T cells and ONC206 against diffuse intrinsic pontine glioma": Official Supplemental Figures.pdf

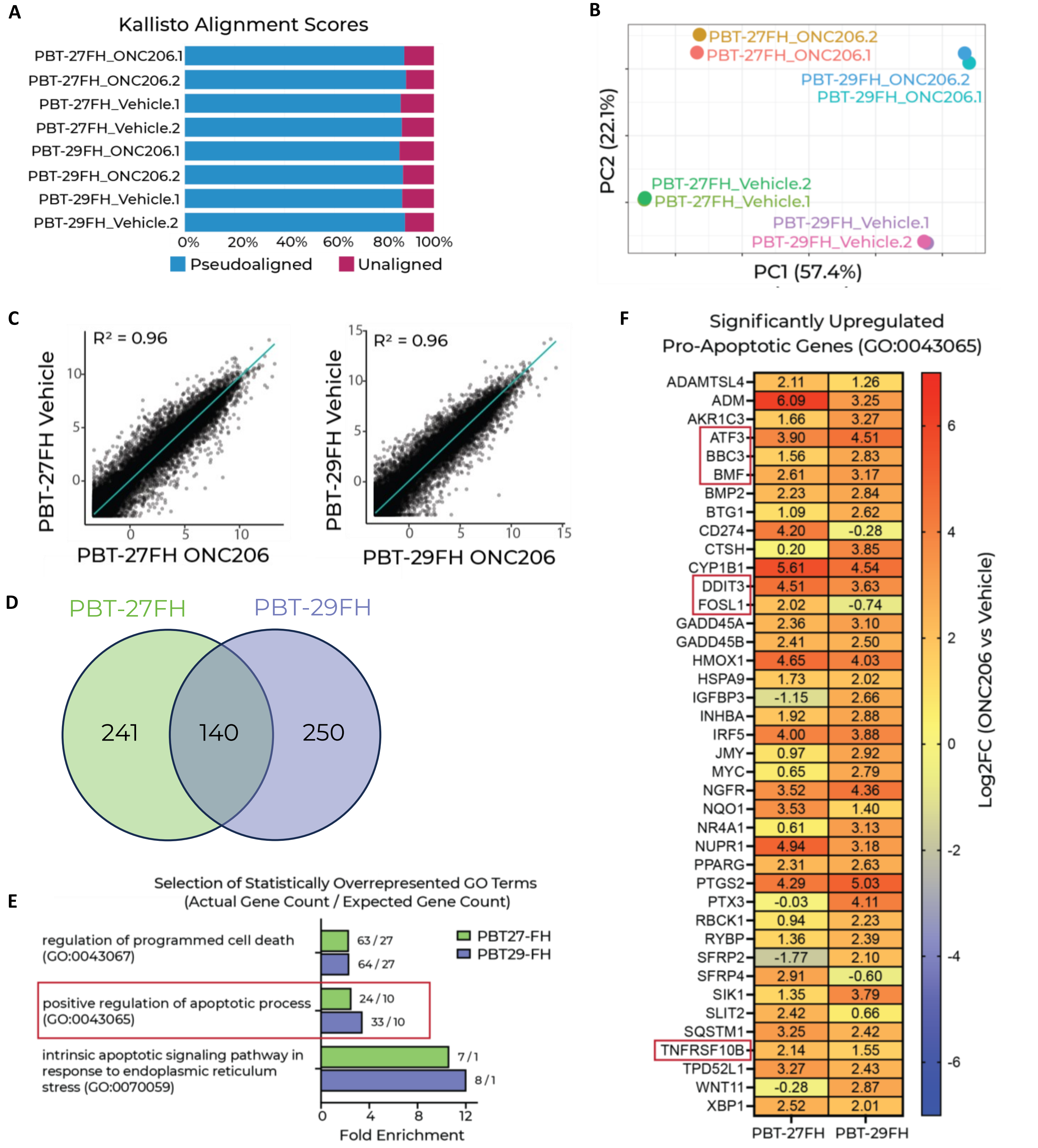


| Gene | PBT-27FH | PBT-29FH |
|---|---|---|
| ADAMTSL4 | 2.11 | 1.26 |
| ADM | 6.09 | 3.25 |
| AKR1C3 | 1.66 | 3.27 |
| ATF3 | 3.90 | 4.51 |
| BBC3 | 1.56 | 2.83 |
| BMF | 2.61 | 3.17 |
| BMP2 | 2.23 | 2.84 |
| BTG1 | 1.09 | 2.62 |
| CD274 | 4.20 | -0.28 |
| CTSH | 0.20 | 3.85 |
| CYP1B1 | 5.61 | 4.54 |
| DDIT3 | 4.51 | 3.63 |
| FOSL1 | 2.02 | -0.74 |
| GADD45A | 2.36 | 3.10 |
| GADD45B | 2.41 | 2.50 |
| HMOX1 | 4.65 | 4.03 |
| HSPA9 | 1.73 | 2.02 |
| IGFBP3 | -1.15 | 2.66 |
| INHBA | 1.92 | 2.88 |
| IRF5 | 4.00 | 3.88 |
| JMY | 0.97 | 2.92 |
| MYC | 0.65 | 2.79 |
| NGFR | 3.52 | 4.36 |
| NQO1 | 3.53 | 1.40 |
| NR4A1 | 0.61 | 3.13 |
| NUPR1 | 4.94 | 3.18 |
| PPARG | 2.31 | 2.63 |
| PTGS2 | 4.29 | 5.03 |
| PTX3 | -0.03 | 4.11 |
| RBCK1 | 0.94 | 2.23 |
| RYBP | 1.36 | 2.39 |
| SFRP2 | -1.77 | 2.10 |
| SFRP4 | 2.91 | -0.60 |
| SIK1 | 1.35 | 3.79 |
| SLIT2 | 2.42 | 0.66 |
| SQSTM1 | 3.25 | 2.42 |
| TNFRSF10B | 2.14 | 1.55 |
| TPD52L1 | 3.27 | 2.43 |
| WNT11 | -0.28 | 2.87 |
| XBP1 | 2.52 | 2.01 |

**Supplemental Figure 1 - ONC206 treatment induces transcriptional reprogramming and upregulation of pro-apoptotic gene signatures in DIPG models.** A. Kallisto alignment scores of bulk RNA-seq reads across all samples (PBT-27FH and PBT-29FH treated with ONC206 or vehicle) indicated high alignment efficiency to the human transcriptome. B. Principal component analysis (PCA) of normalized transcriptomes revealed distinct clustering of samples by treatment condition (ONC206 vs vehicle), indicating strong transcriptional differences. C. Whole transcriptome comparison between treatment conditions (ONC206 vs vehicle) for both cell lines (PBT-27FH and PBT29-FH) depicting best fit line and coefficient of determination (one dot = one gene). D. Venn diagram showing the overlap of significantly upregulated genes (Log2FC > 2, CPM > 1, FDR < 0.05) between the two DIPG models. E. A selection of statistically overrepresented gene ontology (GO) pathways as determined by GO enrichment analysis of significantly upregulated genes across models. F. Heatmap depicting Log2FC values of pro-apoptotic genes (GO: 0043065) significantly upregulated following ONC206 treatment compared to vehicle across DIPG models; key genes such as ATF3, BBC3, BMF, FOSL1, and TNFRSF10B (DR5) are highlighted.

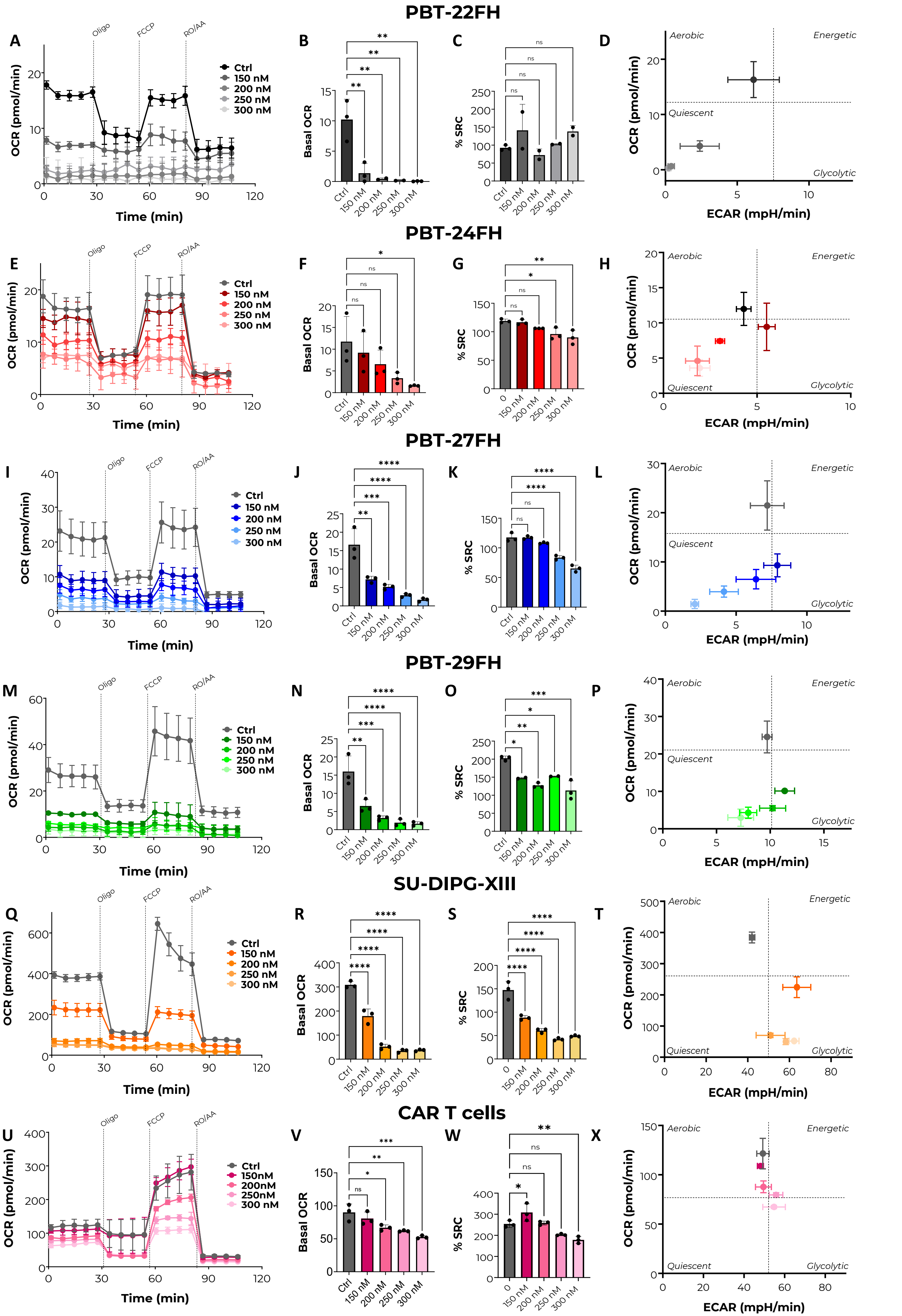
PBT-22FH
A
B
C
D
Oligo
FCCP
RO/AA
Ctrl
150 nM
200 nM
250 nM
300 nM
OCR (pmol/min)
Time (min)
Basal OCR
% SRC
Aerobic
Energetic
Quiescent
Glycolytic
ECAR (mpH/min)
PBT-24FH
E
F
G
H
PBT-27FH
I
J
K
L
PBT-29FH
M
N
O
P
SU-DIPG-XIII
Q
R
S
T
CAR T cells
U
V
W
X
150nM
200nM
250nM
300 nM

**Supplemental Figure 2. ONC206 impairs mitochondrial respiration and promotes metabolic reprogramming in DIPG cells. A–C, E–G, I–K, M–O, Q–S.** Seahorse XF Mito Stress Tests were performed on five DIPG models (PBT-22FH, PBT-24FH, PBT-27FH, PBT-29FH, SU-DIPG-XIII) treated with increasing concentrations of ONC206 (0–350 nM) for 5 days. Oxygen consumption rate (OCR) traces **(A, E, I, M, Q),** basal OCR quantification **(B, F, J, N, R),** and spare respiratory capacity (SRC) **(C, G, K, O, S)** demonstrate a dose-dependent reduction in mitochondrial respiration, with the strongest effects observed at 250–350 nM. **D, H, L, P, T.** Energy phenotype plots (OCR vs. ECAR) show a metabolic shift from aerobic to a more glycolytic or quiescent profile in ONC206-treated DIPG cells, indicating compromised oxidative phosphorylation. **U–X.** Seahorse analyses in CAR T cells treated under the same conditions showed no significant changes in OCR, ECAR, or SRC, suggesting ONC206 selectively impairs tumor metabolism without affecting CAR T cell mitochondrial fitness.
